# Empirical Model of Focused Ultrasound-mediated Treatment for Chemotherapy Delivery to Brain Tumors

**DOI:** 10.1101/2025.01.31.635884

**Authors:** Mohammad Zoofaghari, Martin Damrath, Mladen Veletic, Ilangko Balasingham

## Abstract

Focused ultrasound (FUS) has emerged as a transformative technique for enhancing drug delivery to brain tumors by temporarily and locally disrupting the blood-brain barrier (BBB). Despite significant progress in both preclinical and clinical research, a major challenge remains: the absence of a model that connects the properties of drug particles and FUS sonication parameters to therapeutic effectiveness. In this study, we introduce a novel empirical model that integrates key factors, including drug pharmacodynamics, microbubble kinetics for BBB disruption, intra-brain ultrasound signal propagation, and skull thickness variations. The model defines a new sonication parameter that encapsulates ultrasound signal characteristics and predicts the concentration of therapeutic agents internalized or bound to DNA with an accuracy exceeding 82%. By employing data from previous preclinical studies, this model facilitates the development of precise sonication protocols tailored for clinical applications. These advancements represent a significant step toward personalized FUS-mediated treatments, bridging the gap between experimental research and patient-centered therapies.

## 1. Introduction

Central nervous system (CNS) diseases, including Alzheimer’s, Parkinson’s, multiple sclerosis (MS), brain tumors, stroke, and epilepsy, pose a significant global health burden, affecting millions of people worldwide [1]. For example, around 700,000 people are living with primary brain tumors in the US alone. These conditions are leading causes of death and disability, emphasizing the urgent need for innovative treatments and improved healthcare strategies.

One of the primary challenges in treating CNS diseases is the blood-brain barrier (BBB). The BBB is formed by endothelial cells in the brain’s capillaries protecting the brain from potentially harmful substances in the bloodstream while allowing essential nutrients to pass through. It effectively limits the delivery of chemotherapeutic agents to the tumor site. As a promising strategy to overcome this obstacle, MicroBubble-mediated Focused UltraSound (MB-FUS) has emerged in the last two decades [2]. Ultrasound-mediated therapies utilize the unique properties of ultrasound, such as its non-invasive tissue penetration and capacity to interact with biological structures. Ultrasound signals are produced by ultrasound transducers. Implantable transducers in single-element and multielement devices are used as the ultrasound source as well [3]. The backscattering of ultrasound waves from MBs, generates mechanical disturbances that can temporarily disrupt the BBB, facilitating the passage of administered therapeutic agents through the vascular network to reach target tissues. This effect is enhanced when MBs oscillate in resonance with the ultrasound signal, referred to as *cavitation*. Cavitation can be classified in terms of linearity. In linear cavitation mode, the backscattered signal retains the same frequency as the original ultrasound signal emitted by the transducer. Linear cavitation typically occurs at a low mechanical index (MI ≤0.1), where the ultrasound pressure is relatively mild, causing the microbubbles to oscillate without significant deformation. When the ultrasound pressure increases, the microbubbles oscillate nonlinearly, producing backscattered signals that contain multiple harmonics (higher frequencies) of the original ultrasound signal. This indicates a more complex oscillation pattern of the microbubbles. At very high mechanical indexes, the microbubbles can undergo *inertial cavitation*, characterized by a violent collapse that generates a broadband signal. While this collapse can induce strong mechanical effects and enhance drug delivery, it may also pose a risk of damaging surrounding tissues due to the intense mechanical forces involved.

Studies have demonstrated that ultrasound exposure, in combination with chemotherapeutic agents, can significantly improve the treatment efficacy. For example, for Gliomas, as the most common primary tumors of the brain, an increase in therapeutic effectiveness of up to 3.5-fold and potential survival rate improvements by as much as 30% are reported [4]. However, the effective clinical translation of MB-FUS is hampered by several challenges, including determining the optimal sonication parameters, biological variability, and overcoming translational hurdles [5]. The BBB opening intensity and the pharmacokinetics of therapeutic agents are influenced by specific ultrasound signal characteristics, including acoustic pressure, burst length, and pulse repetition frequency and exposure time [5]. Also, the drug dosage and timing of administration are influential in the success of an MB-FUS-based treatment. Furthermore, the safety concerns are paramount; excessive mechanical stress from microbubbles could result in adverse outcomes like hemorrhage, edema, or apoptosis in healthy tissues [6].

To date, over 100 preclinical studies have examined BBB disruption and the safety of ultrasound exposure. These studies have assessed BBB opening by evaluating the ability of particles, such as MRI contrast agents Gadolinium Diethylene Triamine Pentaacetic Acid (Gd-DTPA) or Evans Blue (EB), to cross the vessel walls. However, these approaches often lack comprehensive pharmacokinetic-pharmacodynamic (PKPD) analyses. Besides, the ultrasound parametric studies conducted in some of these experiments have not yielded a standardized predictive model for the therapeutic response of FUS-mediated treatments. Moreover, permeability measurements derived from MRI using Gd-DTPA as a contrast agent need adaptation for therapeutic agents of different sizes, which may affect their ability to traverse the BBB. In parallel to these preclinical investigations, several clinical trials (Phase I/II) are currently evaluating FUS-mediated treatments for CNS disorders [7, 8, 9, 10, 11, 12] . However, these studies often lack treatment protocols tailored to individual patients and do not fully capture the potential therapeutic improvements resulting from BBB disruption followed by chemotherapy.

This study aims to develop a comprehensive mathematical framework that correlates sonication parameters with drug pharmacodynamics. We define a novel sonication parameter in the linear cavitation regime addressing various features of ultrasound exposure. It integrates all the ultrasound factors into a unified factor considering the impact of each factor on BBB disruption and drug delivery. Our model predicts drug transport through the vessel wall following BBB disruption and its subsequent uptake by the tumor. This is done by employing ordinary differential equations (ODEs) to describe the dynamics of drug transport through the tumor vasculature and transmembrane transport, including DNA intercalation. Our approach is based on data collected from 22 previous preclinical experiments performing ultrasound parametric studies. Hence, our model could reduce the need for costly and challenging in vivo studies paving a road toward improving current and future clinical trials by enabling more precise targeting of drug delivery and sparing patients from ineffective treatments. As illustrated in Fig. 1, the proposed approach bridges the gap between preclinical and clinical studies by developing a comprehensive model that simultaneously incorporates drug characteristics and skull anatomy. This integrated platform enables personalized optimization of ultrasound application and drug administration, tailored to individual patient needs.

**Figure 1:**
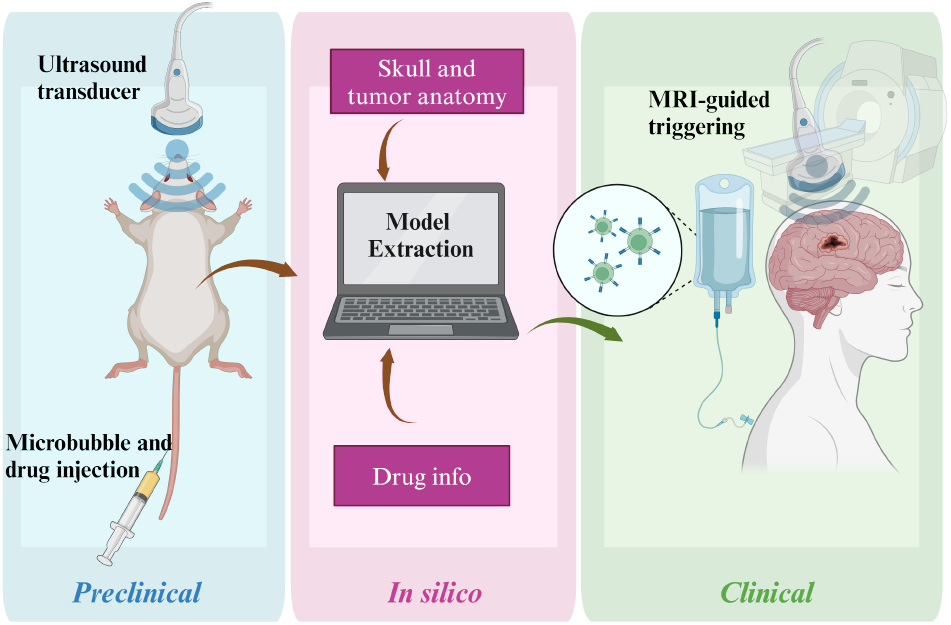
Conceptual illustration of the origin and application of the proposed empirical model. This model utilizes data from previous preclinical studies, along with information on the skull, tumor, and drug, to establish a relationship between therapeutic response and sonication parameters. The model can be used to optimize MRI-guided, focused ultrasound (FUS)-based treatments.

## 2. System model

As previously discussed, FUS-based treatment for CNS diseases depends on ultrasound exposure to facilitate BBB opening and the PK-PD of the administered drug. Key factors in modeling BBB opening include the behavior of the ultrasound signal as it propagates through the skull and brain tissue, along with the dosage and timing of microbubble injection. BBB opening typically occurs within minutes, followed by a closure phase that can last for several hours. Regions with higher vascular density may respond faster due to increased microbubble concentration. The ultrasound signal’s characteristics influence these dynamics, the target locations in the brain and the kinetics of the microbubbles.

### 2.1. The impact of skull anatomy and tumor location

The effects of skull thickness and the ultrasound path from the transducer to the tumor mainly manifest in clinical studies and should be incorporated into the computational model. To tailor the model for individual patients, it is necessary to scale the sonication parameters based on the estimated ultrasonic loss to the target. This propagation loss is characterized by the skull attenuation coefficient α_skl_ and the intracranial path attenuation coefficient α_incl_. Consequently, the loss factor *LF* is given by

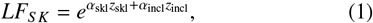

where *z*_skl_ denotes the skull thickness and *z*_incl_ represents the intracranial path length from the transducer to the tumor location. *LF*_*S K*_ can be interpreted as the inverse of *transcranial efficiency* as well. Please note that using an implantable transducer eliminates the intra-skull propagation of ultrasound and significantly reduces the signal attenuation.

### 2.2. The effect of MB administration

As previously discussed, MBs administration is combined with FUS to enhance the effect of ultrasound signals on BBB disruption. The level of the concentration of MBs at the treatment site during the sonication period is a critical factor in determining effective sonication parameters. Here, we focus on the kinetics of MBs and their degradation after administration. Different types of MBs interact with ultrasound in unique ways, depending on their size, oscillatory behavior, and the pressure required to induce effects, which are influenced by shell thickness. However, these specific interactions are not addressed in our model.

Generally, MBs persist for a few minutes, and their concentration can be approximated by an exponential decay described by a first-order ODE. Therefore, we can define MB’s loss factor as

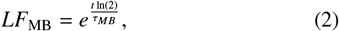

where *t* represents the time after MB injection , and τ_*MB*_ denotes the half-life of the MBs in the bloodstream both in s. The validity of this proposed model will be discussed in the results section.

### 2.3. Proposed sonication parameter

To develop a practical model linking ultrasound signaling to the BBB transvascular rate function, we introduce a novel sonication parameter (*S P*) given by

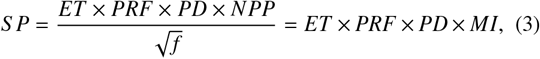

where *ET* is the exposure time in s , *PRF* is the pulse repetition frequency in Hz, *PD* is the pulse duration in ms, *NPP* is the negative peak pressure in MPa, and *f* is the ultrasound frequency in MHz and *MI* denotes the mechanical index measuring the instantaneous cavitation potential. *S P* represents the *cumulative mechanical impact* of ultrasound over the total exposure time. It combines the instantaneous mechanical cavitation potential with the temporal characteristics of the ultrasound signal (exposure time, pulse repetition frequency, and pulse duration). Higher values of *S P* indicate greater total mechanical bioeffect of ultrasound in our application.

In order to incorporate the loss effects as discussed in previous subsections, the *S P* given by the model should be scaled by loss factors *LF*_*S K*_ and *LF*_*MB*_ given by (1) and (2), respectively as

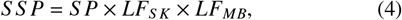

where *S S P* is scaled sonication parameter. Given the limited dataset and the inherent variability due to differing experimental conditions, we aim to establish a linear relationship between *S P* and the therapeutic response. This approach is chosen to mitigate the risk of overfitting. Additionally, a linear model facilitates a matrix-based formulation of the problem, which is advantageous for multi-target scenarios represented as a MIMO system. Such a framework can be extended in future work to design optimized drug delivery strategies across multiple targets.

In order to relate the therapeutic response to the sonication parameter, we first need to discuss the PK-PD aspects of the therapeutic agent.

### 2.4. Pharmacokinetics and pharmacodynamics of drug delivery through BBB

Significant research has been conducted and mathematically implemented on the pharmacodynamics of the BBB and the therapeutic agents [13, 14, 15]. In this study, we focus on a single mechanism as a proof of concept to demonstrate the ability to predict drug delivery mediated by nuclear translocation using the transvascular rate function and interaction model in FUS-based treatment. Nevertheless, equations (7) and (8) can be adapted or replaced with formulations describing the dynamics of other therapeutic agents, such as small molecule inhibitors or monoclonal antibodies targeting specific cell surface receptors or antigens. We use sets of ODEs to describe the drug’s transport from the bloodstream to the cell nucleus, often assuming a single-target pathway as shown below:

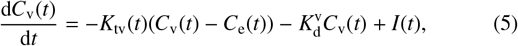

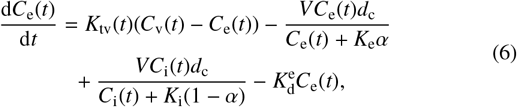

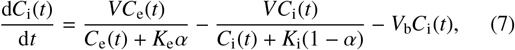

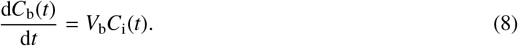

This set of ODEs includes key variables such as concentration parameters *C*_v_(*t*), *C*_e_(*t*), *C*_i_(*t*), and *C*_b_(*t*); kinetic parameters 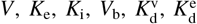; and biological factors *d*_c_, α. The ultrasound-related parameter *K*_tv_(*t*) and drug infusion function *I*(*t*) are also considered. Here, *C*_v_(*t*) represents the vascular agent concentration, *C*_e_(*t*) denotes extracellular agent concentration, *C*_i_(*t*) pertains to internalized agents, and *C*_b_(*t*) indicates the concentration of DNA-bound agents. The rate of transmembrane transport is given by *V*, with *K*_e_ and *K*_i_ serving as the Michaelis constants for this process. *V*_b_ represents the rate at which particles bind to DNA, and is particularly relevant for drugs such as doxorubicin. 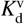 accounts for drug excretion via organs like the kidney and liver, while 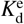 represents the rate of extracellular degradation. The parameter *d*_c_ denotes tumor cell density, and α represents the volume fraction of the extracellular matrix. The parameter *K*_tv_(*t*), which denotes the transvascular rate, can be modulated by ultrasound exposure and is typically modeled as an exponential function:

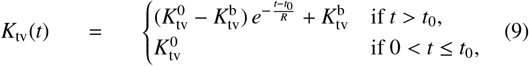

where 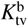 is the baseline transvascular rate, and *t*_0_ denotes the time interval in which the BBB is open with a transvascular rate of 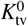. After the sonication process stops, *K*_tv_(*t*) decreases, reflecting the BBB closure with a time constant *R*. This function is generally measured through the extravasation of MRI agents, such as Gd-DTPA, into the tissue. The function also depends on the molecular weight of the administered particle and is scaled by (1 − 0.5 log(*M*_r_)), where *M*_r_ is the molecular weight ratio relative to Gd-DTPA [16]. The BBB closure time constant must also be adjusted according to the therapeutic agent size [17]. To calculate the response of internalized or DNA-bound agents, the system of ODEs (5)-(8) is solved with the input function defined as

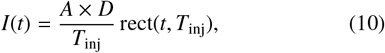

where *A* is the inverse of the volume of distribution in plasma, *D* is the total injected dosage, and rect(*t, T*_inj_) represents a rectangular pulse function with duration *T*_inj_ starting at *t* = 0.

Please note that during the agent’s administration, the intensity of BBB disruption as a result of ultrasound exposure should be tailored to the specific therapeutic agent. Also, the therapeutic window of an agent should be observed in the modeling to manage the drug efficacy, adverse effects and cytotoxicity.

## 3. Numerical results

### 3.1. Empirical model for 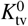 in terms of S P

As discussed in Section 2, the initial transvascular rate constant, 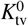, is required to calculate the therapeutic response (e.g., concentration of bound-to-DNA agents). Instead of 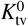, many studies investigate EB staining patterns, traditional signal intensity change (SIC) maps in MRI T1-weighted images (DCE-T1), or Gd-based area-under-curve (Gd-AUC) maps, which provide data for deriving the transvascular rate function. Fortunately, there is a strong correlation between these criteria, which can be used to establish a model for 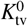. Several studies on the effects of sonication parameters for BBB opening report the *S IC* in DCE-T1 map, allowing us to relate 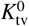 to the sonication parameter. If we use the MRI contrast agent, the signal intensity increase shows the BBB disruption. Please note that the endothelial pores are usually wide enough even without any sonication for the water molecules to cross the BBB. Accordingly, assessment of FUS-opening is carried out using T1-weighted images and inspection of edema and hemorrohage is usually done using the T2-weighted MRI scans.

We use data from [18], as shown in Fig. 2a, with a linear fitting function that relates 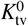 to *S IC*. Using this function, along with the data provided in Table 1 from 22 preclinical studies that demonstrate *S IC* for various ultrasound parameters, we derive a relationship for 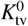 in terms of the sonication parameter (*S P*), as shown in Fig. 2b. To assess the fitting performance, we calculate the R-squared and Root Mean Square Error (RMSE) metrics. R-squared measures the correlation between the actual and estimated values, with values closer to one indicating a better fit. R-squared is given by

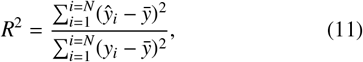

where *y*_*i*_, *ŷ*_*i*_ and 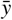 stand for real, estimated, and mean value of data and *N* denotes the dataset size. We also exploit the RMSE as the fit standard error estimatng the standard deviation of the random component in data given by

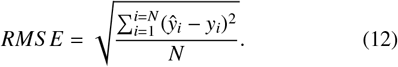

**Table 1:** Ultrasound parameters, calculated sonication parameter and the signal intensity change for 22 preclinical studies included in this paper for model extraction.

| PD[ms] | PRF[Hz] | NPP[MPa] | f [MHz] | ET[s] | SP[Pa s<sup>1.5</sup>] | SIC [%] | Reference |
| --- | --- | --- | --- | --- | --- | --- | --- |
| 1 | 1 | 0.3 | 0.5 | 60 | 25 | 1.8 | [19] |
| 10 | 1 | 0.3 | 0.5 | 30 | 127 | 5.9 | [19] |
| 10 | 1 | 0.3 | 1.6 | 60 | 142 | 5.1 | [19] |
| 10 | 1 | 0.2 | 0.69 | 60 | 144 | 1 | [20] |
| 10 | 1 | 0.2 | 0.5 | 60 | 170 | 1.8 | [19] |
| 10 | 1 | 0.3 | 0.5 | 60 | 255 | 13.7 | [19] |
| 10 | 1 | 0.3 | 0.5 | 60 | 255 | 5 | [19] |
| 10 | 1 | 1.1 | 1.5 | 30 | 269 | 12 | [21] |
| 10 | 1 | 0.6 | 1.6 | 60 | 284 | 9 | [19] |
| 10 | 1 | 0.4 | 0.69 | 60 | 288 | 1.8 | [20] |
| 10 | 1 | 0.5 | 0.69 | 60 | 361 | 7 | [20] |
| 10 | 1 | 0.3 | 0.4 | 90 | 369 | 22 | [18] |
| 10 | 1 | 1.9 | 1.5 | 30 | 465 | 12 | [21] |
| 10 | 1 | 0.3 | 0.5 | 120 | 509 | 23.1 | [19] |
| 10 | 1 | 0.6 | 0.5 | 60 | 509 | 21 | [19] |
| 10 | 2 | 0.3 | 0.5 | 60 | 509 | 13.7 | [19] |
| 10 | 1 | 0.8 | 0.69 | 60 | 577 | 18 | [20] |
| 10 | 1 | 0.54 | 1.18 | 120 | 594 | 26 | [22] |
| 10 | 1 | 2.54 | 1.5 | 30 | 600 | 16 | [21] |
| 10 | 1 | 1.1 | 0.69 | 60 | 794 | 20 | [20] |
| 10 | 1 | 3.5 | 1.5 | 30 | 857 | 30 | [21] |
| 10 | 1 | 1.5 | 0.69 | 60 | 1083 | 32 | [20] |
| 10 | 5 | 0.3 | 0.5 | 60 | 1272 | 25 | [19] |

**Figure 2:**
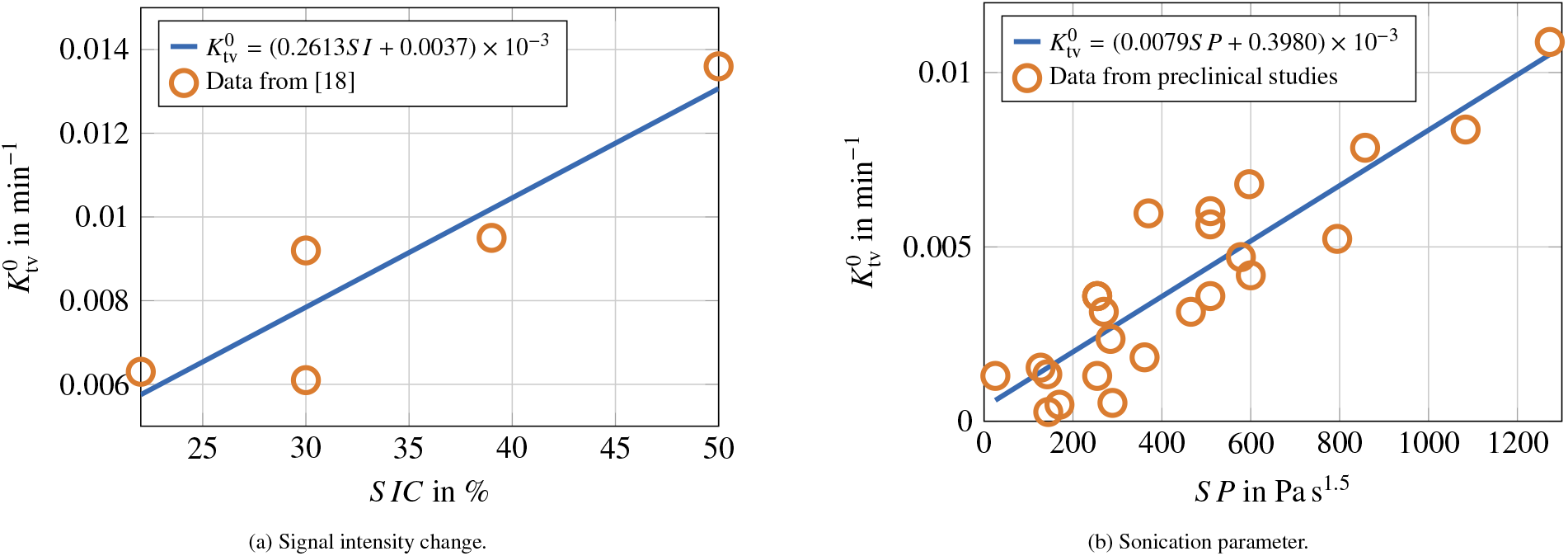
Initial transvascular rate constant versus (a) signal intensity change in MRI T1-weighted images given by [18] and (b) sonication parameter for 22 preclinical studies, as presented in Table 1, along with fitting model.

Fig. 2b shows the linear fitting function relating the 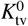 to the sonication parameter with R-squre= 0.8 and RMSE= 0.0011.

### 3.2. Evalution of the proposed model for 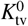 by the clinical data

There is a notable scarcity of studies addressing transcranial characteristics and the transvascular rate coefficient 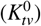 across different ultrasound parameters and patient profiles. To evaluate the accuracy of our model for clinical applications, we utilized data from [12]. Table 2 presents data about transcranial efficiency and the measured 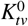 values for four patients. The final two columns of the table provide the 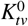 values estimated by our model alongside the corresponding estimation errors relative to the measured data. Despite significant physiological differences between the mouse model and the human brain, the model demonstrates reasonable accuracy in estimating the rate coefficient. It is worth noting that the estimation errors of the model are lower than the measurement errors reported in clinical data, highlighting its potential reliability for practical applications.

**Table 2:**
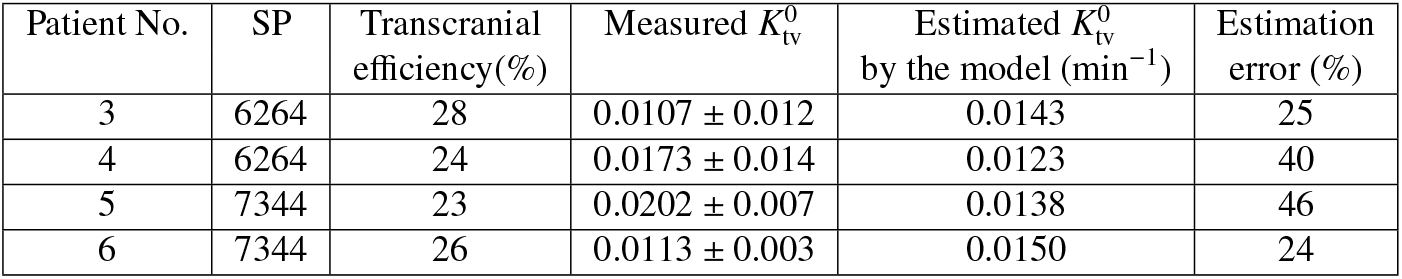
Estimation error of the proposed model for the clinical measurement data reported by [12].

### 3.3. Empirical model for therapeutic response in terms of S P

In this subsection, we derive a model for the therapeutic response of the drug, building on the previously extracted model for 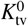. The therapeutic response can be defined based on the drug’s functionality with target cells, involving *C*_*i*_(*t*) or *C*_*b*_(*t*). Fig. 3a and Fig. 3b show the obtained data points for *C*_*i*_(∞) and *C*_*b*_( ), along with the linearly fitted functions for the 22 preclinical studies, with rate parameters provided in Table 1. The R-squared and RMSE values for Fig. 3a are 82% and 6.5 × 10^−19^, and for Fig. 3b are 83% and 2.2 × 10^−14^, respectively. These parameters demonstrate that the linear fitting accurately describes the therapeutic response behavior versus the sonication parameters.

**Figure 3:**
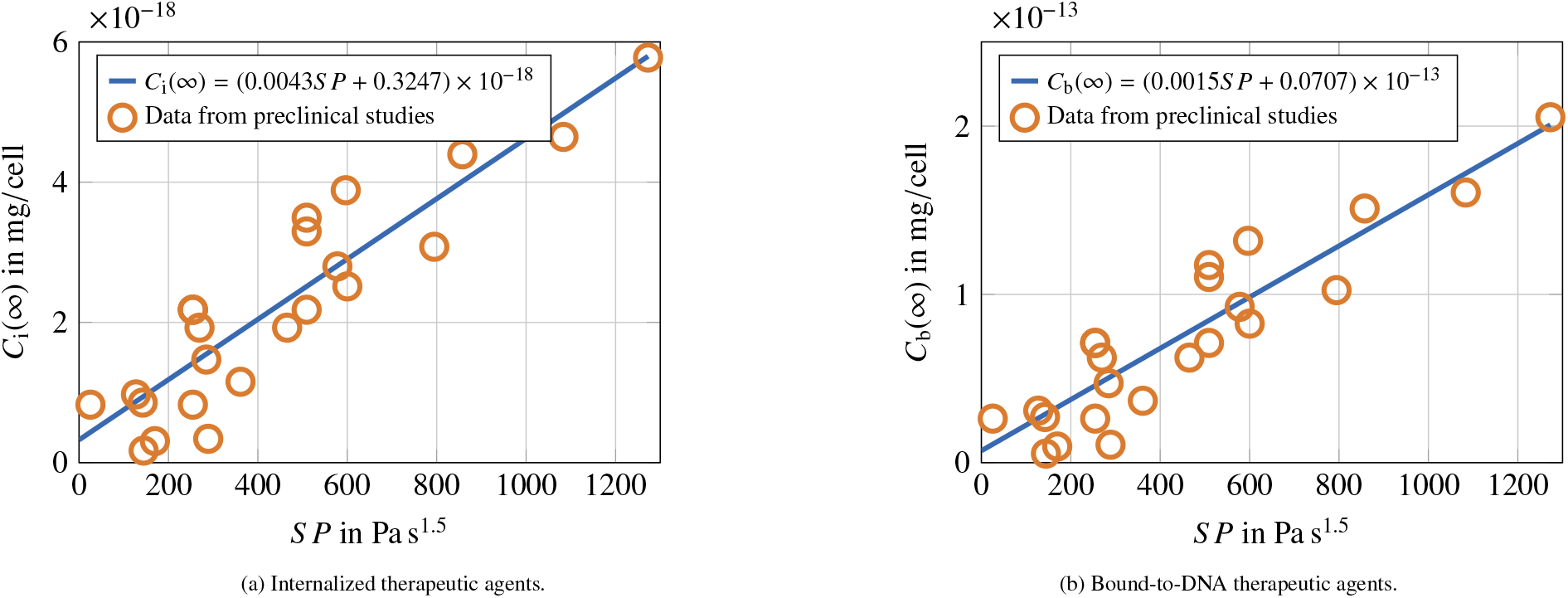
Concentration of (a) internalized therapeutic agents and (b) bound-to-DNA therapeutic agents derived from agent pharmacokinetics ODEs and the transvascular rate constant model, based on 22 preclinical datasets of signal intensity change, along with the fitted linear function versus sonication parameter.

The estimation of the binding rate constant *V*_*b*_ is subject to inherent uncertainty, as values reported in the literature may vary widely, potentially leading to significant overestimation or underestimation. Fig. 4 illustrates the proposed model function (solid line) as well as the variation in the concentration of bound-to-DNA agents, *C*_*b*_(∞), across a range of 10-fold underestimation (circle denoting the lowest *C*_*b*_(∞) for each *S P*) to 10-fold overestimation (circle denoting the highest *C*_*b*_(∞) for each *S P*) of *V*_*b*_ for each preclinical dataset. It is evident that the variation in *C*_*b*_(∞) becomes more pronounced under higher sonication parameters, where the accuracy of the proposed model diminishes. Moreover, overestimation of *V*_*b*_ can compromise the model’s validity more than underestimation of *V*_*b*_ . To assess the sensitivity of our model to changes in *V*_*b*_, we plotted the R-square and RMSE values against the logarithm of the fold change in *V*_*b*_. As shown in Fig. 5, the model achieves a satisfactory goodness of fit for fold changes greater than −0.4.

**Figure 4:**
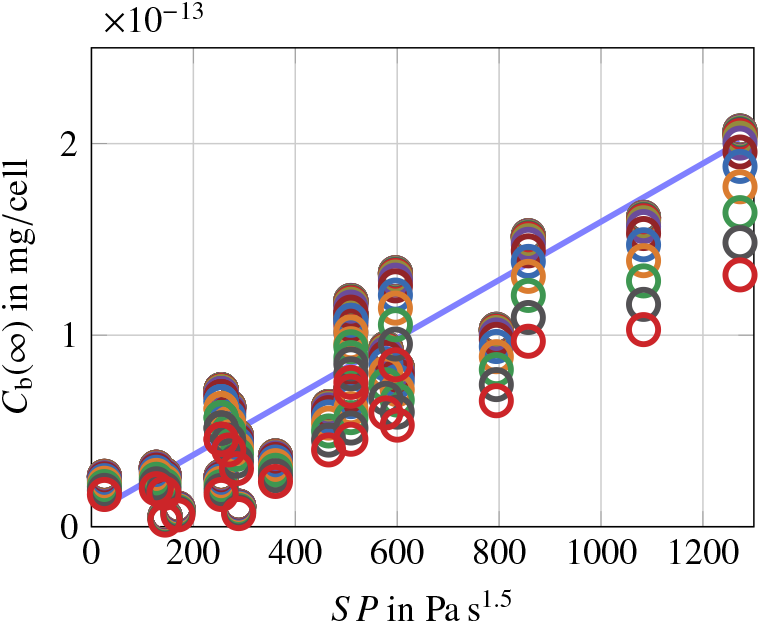
Variation of *C*_*b*_(∞ ) with respect to 10-fold changes in *V*_*b*_ for each preclinical dataset.

**Figure 5:**
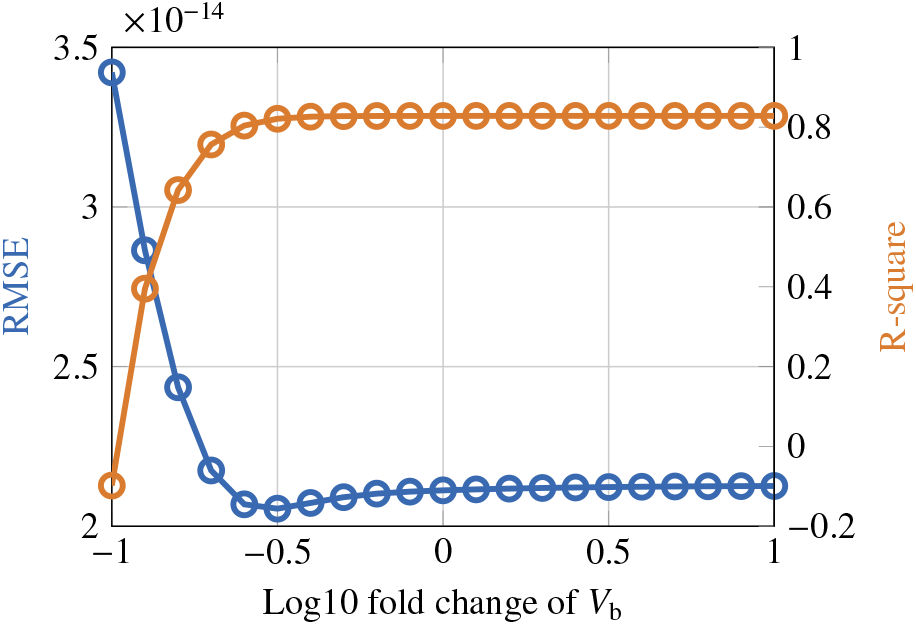
R-square values as a function of the logarithm of the fold change in *V*_*b*_.

**Table 3:**
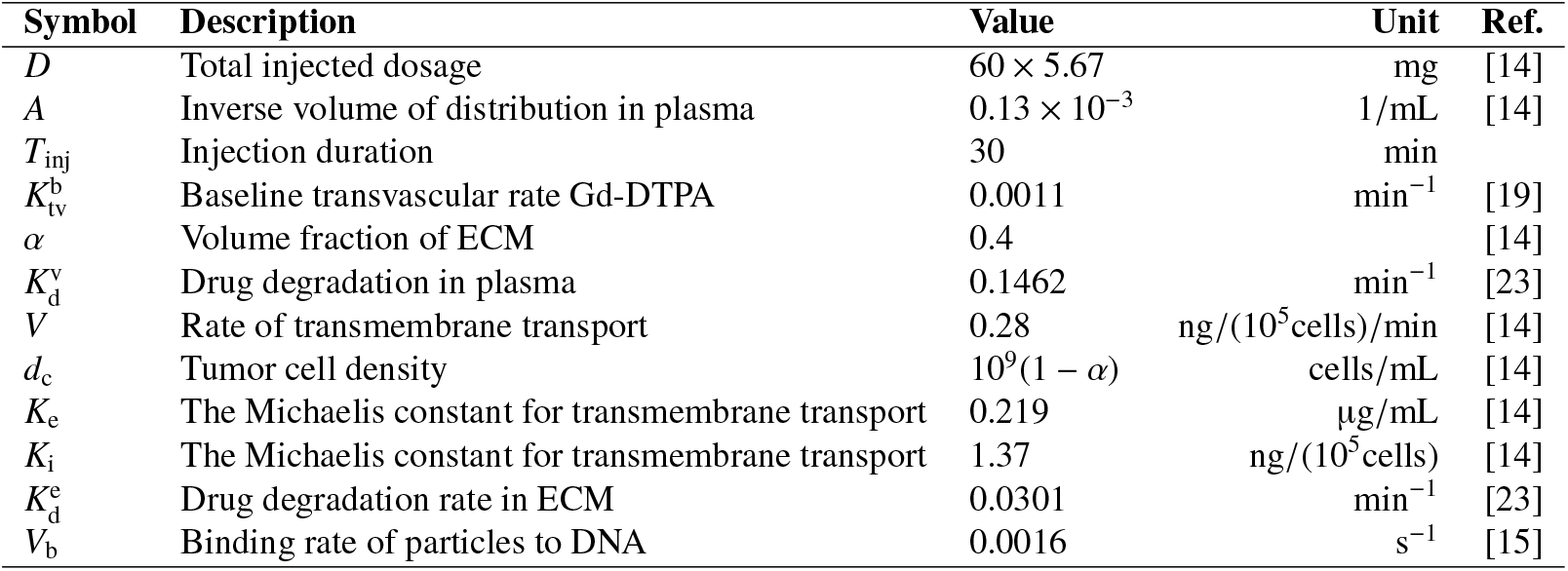
Model parameters and their units and values considered in this paper.

### 3.4. Impact of skull topography

As previously mentioned, intracranial propagation can significantly attenuate the ultrasound signal, primarily due to skull thickness. This requires an external sonication scheme that is adjusted using *LF*_*S K*_, as defined in (1), to ensure the same ultrasound effect at the tumor site as in the absence of intracranial attenuation. Ignoring the path loss from the skull to the tumor, Fig. 6a depicts the *C*_*b*_(∞) model for three different skull thicknesses. As shown, the slope of the linear model describing *C*_*b*_(∞) in terms of SP decreases as skull thickness increases. This indicates that in a clinical study, increased sonication is necessary for patients with thicker skulls to achieve the same therapeutic response as those with thinner skulls.

**Figure 6:**
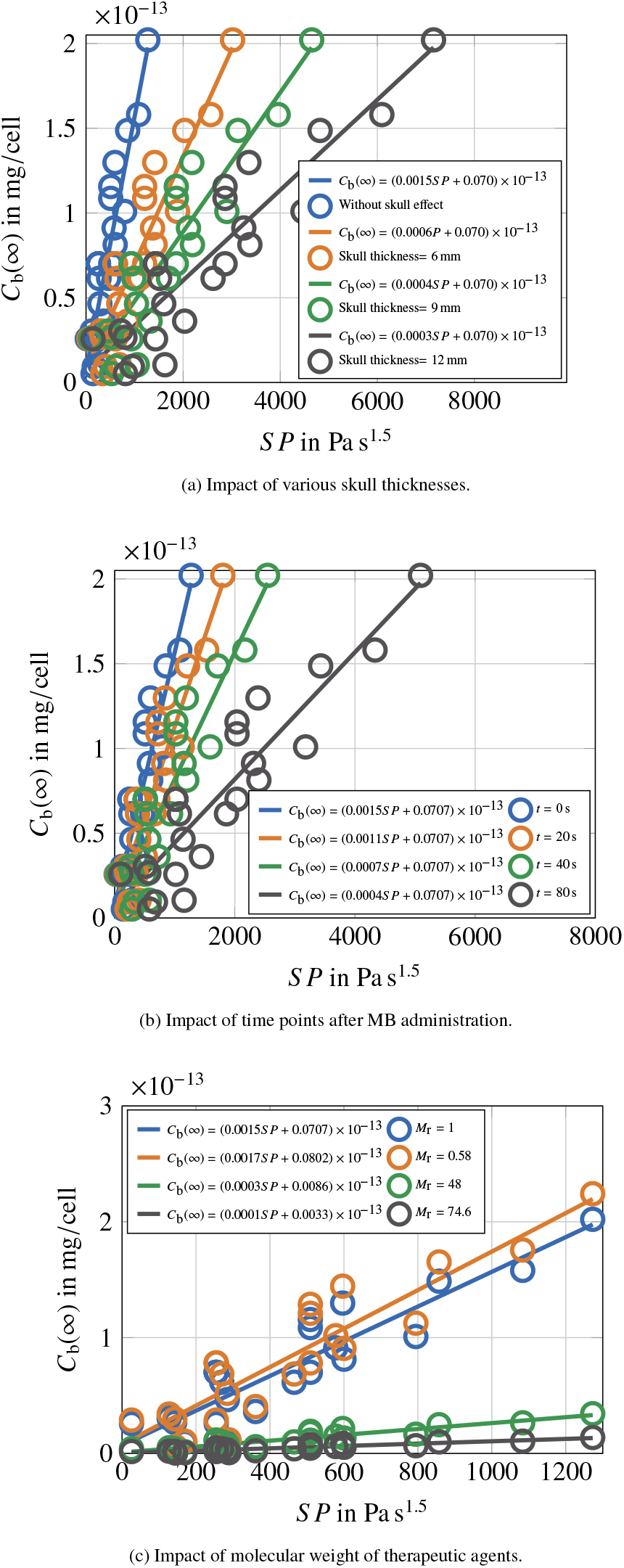
Therapeutic response models for various (a) skull thicknesses, (b) time points after MB administration, and (c) relative molecular weight of therapeutic agents.

### 3.5. Validation of the model for LF_MB_ and impact of MB dosage on therapeutic response

As discussed in Sec. 2.2, the kinetics of microbubbles (MB) can be related to the sonication parameters by an exponential sonication factor given by (2). Here, we utilized the preclinical data provided by [24] to evaluate this equation. The data were collected from the cavitation dosage reported in Fig. 4 of [24] for 9 spots serially sonicated in a mice study. In this study, each spot is exposed to ultrasound for 20 seconds, which determines the time step of the data points. The normalized cavitation data and the fitted function versus time are shown in Fig. 7. It is observed that the shown fitted exponential function closely matches the model provided by (2), where τ_*MB*_ = 40 sec, which is consistent with values reported in the literature for utilized MBs [25].

**Figure 7:**
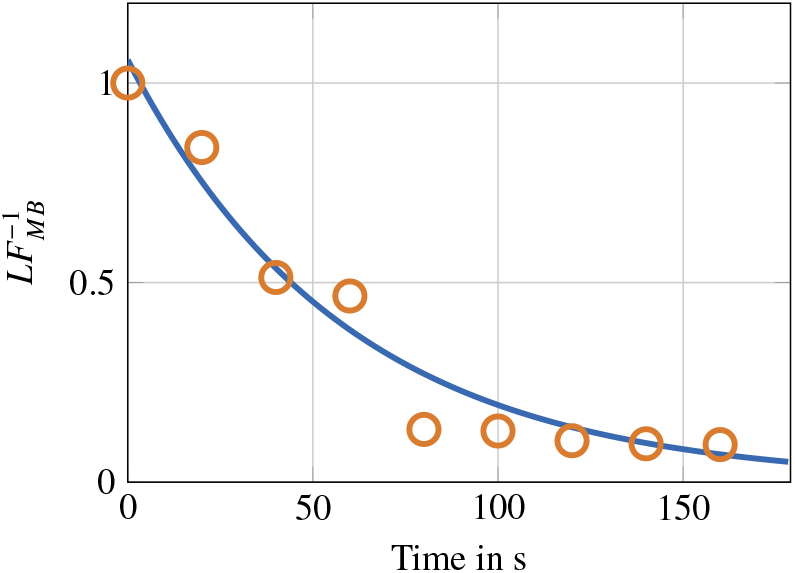
Inverse of loss factor due to MB degradation approximated by the normalized cavitation data given by [24](red circles). The solid line denotes the fitting function.

Fig. 6b illustrates the therapeutic response models as a function of *S P* at different time points *t* following MB administration. This considers *LF*_MB_ given by (2) and the half-life estimated by the fitting exponential function in Fig. 7. As observed, an increase in sonication intensity is required over time due to the decreasing concentration of MBs in the bloodstream. This observation underscores the critical importance of precise scheduling in FUS-based treatments, especially in scenarios involving multiple target regions.

### 3.6. Impact of relative molecular weight of therapeutic agents

As mentioned previously, the therapeutic response of drug agents can vary significantly depending on their molecular weight. This factor must be considered when designing a FUS-based therapy, as larger particles require more intense sonication to achieve the same therapeutic response as smaller particles. To address this issue, we consider three different drug types that vary considerably in size to evaluate the performance of our model. Fig. 6c illustrates the extracted models for different therapeutic agents: free doxorubicin (*M*_*r*_ = 0.58), polymeric nanoparticles (*M*_*r*_ = 48), liposomal doxorubicin (*M*_*r*_ = 74.6), and Gd-DTPA as a reference in our studies [26]. As shown, the therapeutic response of liposomal doxorubicin is approximately 17 times lower than that of free doxorubicin, given the same sonication parameters. This demonstrates the necessity of adjusting the sonication scheme for liposomal doxorubicin and even the injection dosage, as there are limitations on ultrasound intensity to avoid tissue damage.

## 4. Conclusion

In this study, we developed a comprehensive model for predicting the therapeutic response of focused ultrasound (FUS)-based treatments as a function of the sonication parameter (SP), specifically targeting brain tumors . The sonication parameter is introduced as a quantitative measure that characterizes the ultrasound signal, which can be adjusted by clinicians to modulate drug concentration at specific target sites within the tumor. Our model integrates key factors, including intracranial ultrasound attenuation, drug agents size, and the intravascular pharmacokinetics of MBs. The results demonstrate that the proposed model is robust to variations in reaction rates, which is critical for accommodating the inherent biological variability between patients with brain tumors .

The model presented in this study focuses primarily on the parameters of FUS-mediated treatments associated with the sonication protocol. However, important factors such as the vascular density, vessel diameter, and overall configuration of the targeted region have not been included. Addressing these aspects would require extensive preclinical datasets with sufficient diversity in vascular characteristics. This limitation underscores an opportunity for future research aimed at enhancing the model to incorporate these biological complexities. Also, future work may focus on validating this model through clinical trials, aiming to confirm its predictive accuracy and clinical utility in brain tumors treatment. Clinical validation of the drug delivery model for the FUS-based treatment presented in this study could be performed using biopsy pathology tests. This approach would involve at least two tests using distinct sonication parameters to assess the fold change in drug concentration within the tumor tissue. The observed fold change would then be compared to the slope predicted by the model. To enhance validation, the biopsy results could be complemented with imaging techniques or other biomarkers that indicate drug distribution and concentration in the tissue.Additionally, we plan to integrate this model with optimization algorithms to maximize the therapeutic efficacy of FUS-based treatments, particularly in multi-targeted treatment scenarios where precise control of drug delivery is critical.

## Conflict of Interest

The authors declare that they have no known competing financial interests or personal relationships that could have appeared to influence the work reported in this paper.

## Data Availability Statement

The data supporting the findings of this study are available upon the reasonable request of corresponding author.

## Notes

### Competing Interest Statement

The authors have declared no competing interest.

